# Cytosolic N6AMT1-dependent translation supports mitochondrial RNA processing

**DOI:** 10.1101/2024.07.02.601698

**Authors:** Mads M. Foged, Emeline Recazens, Sylvain Chollet, Miriam Lisci, George E. Allen, Boris Zinshteyn, Doha Boutguetait, Christian Münch, Vamsi K. Mootha, Alexis A. Jourdain

## Abstract

Mitochondrial biogenesis relies on both the nuclear and mitochondrial genomes, and imbalance in their expression can lead to inborn error of metabolism, inflammation, and aging. Here, we investigate N6AMT1, a nucleo-cytosolic methyltransferase that exhibits genetic co-dependency with mitochondria. We determine transcriptional and translational profiles of N6AMT1 and report that it is required for the cytosolic translation of TRMT10C (MRPP1) and PRORP (MRPP3), two subunits of the mitochondrial RNAse P enzyme. In the absence of *N6AMT1*, or when its catalytic activity is abolished, RNA processing within mitochondria is impaired, leading to the accumulation of unprocessed and double-stranded RNA, thus preventing mitochondrial protein synthesis and oxidative phosphorylation. Our work sheds light on the function of *N6AMT1* in protein synthesis and highlights a cytosolic program required for proper mitochondrial biogenesis.

## Introduction

Most of the mitochondrial proteome is encoded in the nucleus, and only 13 mitochondrial proteins, all of which are required for mitochondrial oxidative phosphorylation (OXPHOS), are derived from the mitochondrial genome (mt-DNA). To ensure the assembly of the respiratory chain complexes and respond to environmental changes, the expression of both genomes must be tightly coordinated. For example, transcriptional mechanisms such as those relying on PGC-1α (peroxisome proliferator-activated receptor gamma coactivator 1-alpha) and NRF1 (nuclear respiratory factor 1), two master regulators of mitochondrial biogenesis, can promote mt-DNA expression by inducing transcription of *TFAM*^1,2^, a nuclear-encoded rate-limiting factor in mt-DNA replication and transcription. In addition, CLUH (clustered mitochondria protein homologue) and 4E-BPs (eukaryotic translation initiation factor 4E-binding proteins), cytosolic proteins which respond to nutrient deprivation, promote the translation of nuclear-encoded mitochondrial proteins^3,4^.

Within mitochondria, both strands of the mt-DNA molecule are transcribed as complementary polycistronic transcripts containing rRNAs, tRNAs, mRNAs, and non-coding RNAs (ncRNAs). Newly synthesized mt-RNA accumulates in mitochondrial RNA granules (MRGs)^5–7^ where the mitochondrial protein-only RNAse P complex (composed of TRMT10C/MRPP1, HSD17B10/MRPP2 and PRORP/MRPP3/KIAA0391) and RNAse Z (ELAC2) process polycistronic mt-RNA by excising tRNAs according to the tRNA punctuation model^8,9^. Blocking mitochondrial RNA processing leads to the retention of newly transcribed mt-RNA in MRGs^5^. An integral part of RNA processing is the degradation of ncRNAs by the mitochondrial RNA degradosome (SUPV3L1, PNPT1), which fine-tunes mitochondrial gene expression and prevents accumulation of the double-stranded RNA (dsRNA) generated through complementarity between the bidirectional mt-RNA transcripts^10,11^. Inborn errors in the machinery of mitochondrial RNA degradation result in mt-dsRNA accumulation, leakage into the cytoplasm, and a chronic interferon response in humans^11^.

In two recent genetic screens, we and others have highlighted a possible role for the nucleo-cytoplasmic multi-substrate methyltransferase N6AMT1 in OXPHOS^12,13^. N6AMT1 (also known as HEMK2, KMT9) was originally identified as a putative N_6_-adenine-specific DNA methyltransferase, but the existence of N6-methyladenosine in human DNA is uncertain^14,15^, and recent *in vitro* assays as well as structural studies have questioned its DNA methyltransferase activity^16,17^. In bacteria, yeast, and higher eukaryotes, N6AMT1 and its homologues (prmC, Mtq2) methylate translation release factors (RFs) on a highly conserved GGQ motif^18–22^. In bacteria, RF1/RF3 methylation increases the speed and accuracy of translation termination^23^, while the role of eRF1 methylation in eukaryotes is less clear^21,24,25^. In yeast, Mtq2 interacts with proteins of the 60S cytosolic ribosomal subunit, and an *MTQ2*-null strain exhibits defects in ribosome biogenesis, with no defects in translation termination^24^. In *Drosophila melanogaster*, HemK2 depletion leads to ribosome stalling and a decline in mRNA stability^26^. In mice, genetic ablation of *N6amt1* is lethal^25^, but to date, its impact on mammalian translation has not been investigated. Additional protein methylation substrates have also been identified for N6AMT1, including histone H4, which undergoes lysine 12 monomethylation affecting transcription of cell cycle and cancer-related genes^27,28^, as well as methylation of translation-related factors^44,45^. N6AMT1 may also modulate arsenic-induced toxicity, possibly by converting arsenic derivatives into less toxic forms, and polymorphisms in *N6AMT1* are associated with arsenic resistance in Andean women^29,30^. It is currently unclear which of the multiple roles attributed to *N6AMT1* accounts for its putative mitochondrial phenotype.

Here, we investigate *N6AMT1* and its role in mitochondrial function. We report that *N6AMT1* is required for the cytosolic synthesis of two key subunits of the mitochondrial RNA processing machinery. Depletion or catalytic inactivation of *N6AMT1* results in the accumulation of unprocessed and double-stranded mt-RNA, leading to decreased mitochondrial protein synthesis, progressive collapse in the mitochondrial respiratory chain, and an interferon response. Our work identifies a novel mechanism required for the coordinated expression of the nuclear and mitochondrial genomes based on *N6AMT1*.

## Results

### N6AMT1 is a nucleo-cytosolic protein required for mitochondrial gene expression

To improve our understanding of *N6AMT1* in humans in the context of its previously reported functions, we analyzed the genetic dependency on *N6AMT1* across 1,100 cancer cell lines from the Cancer Cell Line Encyclopedia (CCLE)^31^. We found that cells are selectively dependent on *N6AMT1* (CCLE Chronos score < -0.5 in 35.3% of the cell lines), irrespective of their lineages (Figures 1A and S1A). Comparing the *N6AMT1*-dependency profile across the 1,100 cancer cell lines with the dependency profiles of 15,677 non-essential genes, we found that *N6AMT1* correlated best with nuclear-encoded genes coding for mitochondrial proteins (MitoCarta3.0 genes^32^) (Figures 1B-C and S1B, Supplementary tables S1-S2). This observation supports a possible role for *N6AMT1* in mitochondrial function; however, consistent with previous reports^33,34^, we found no colocalization of N6AMT1 with mitochondria using confocal microscopy (Figure 1D).

**Figure 1.**
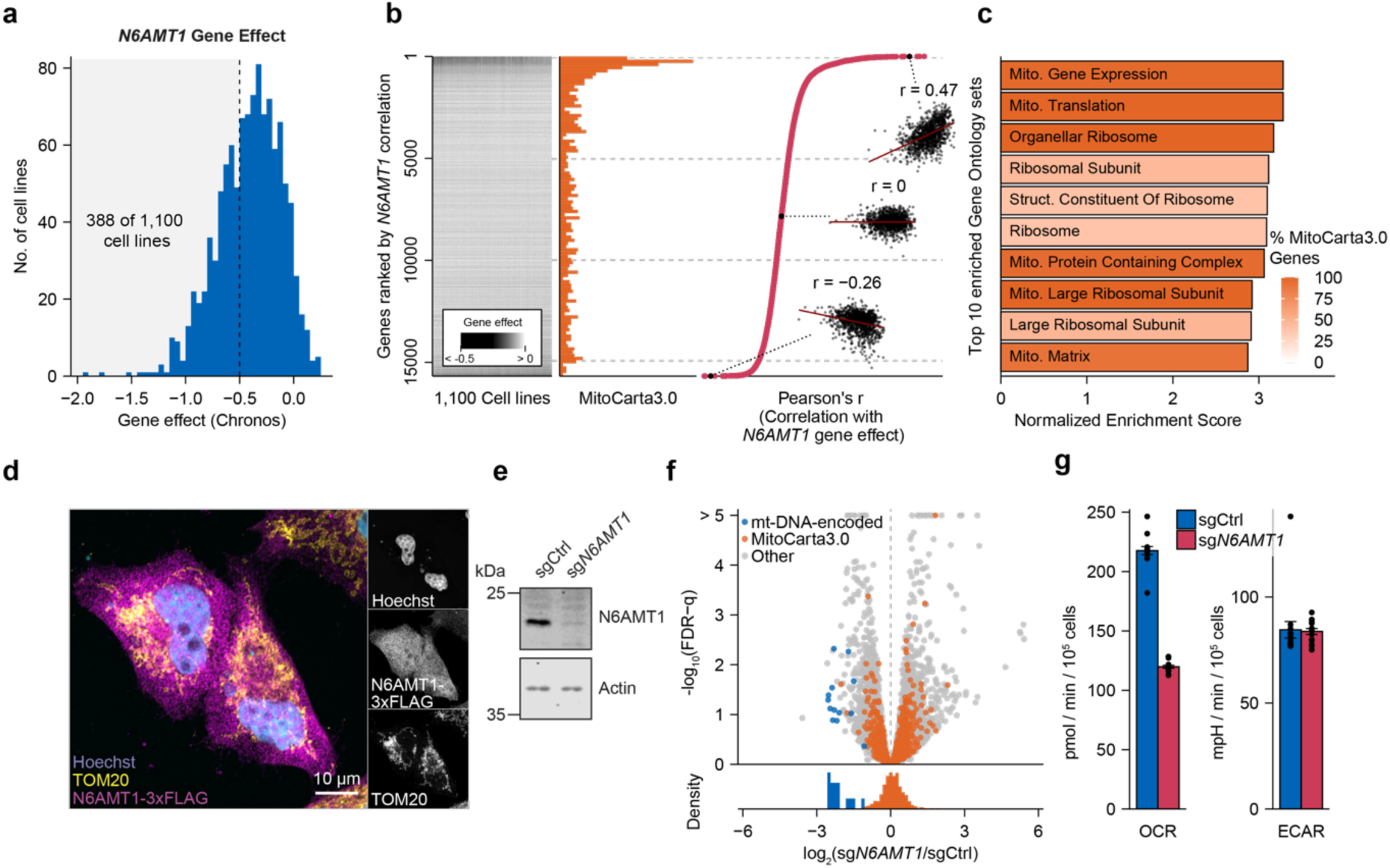
Mitochondrial impact of *N6AMT1* is independent of nuclear transcription. **a** Cell line distribution of *N6AMT1* Chronos scores across 1,100 cell lines of the Cancer Cell Line Encyclopedia (CCLE). Density represents the relative number of cells lines with an *N6AMT1*-gene effect score within the range of each bin. Shaded area represents highly *N6AMT1*-dependent cell lines (388 of 1,100 cell lines with Chronos score < -0.5). **b.** Analysis of *N6AMT1* gene effect correlations. Gene-effect scores (Chronos scores; left panel) of 15,677 non-essential genes in 1,100 cell lines from the CCLE were correlated with *N6AMT1* gene effects and ranked according to Pearson correlation coefficients (right panel). The distribution of MitoCarta3.0 genes is indicated in the middle panel. Inserts show example correlations between *N6AMT1* and three arbitrarily selected genes. Pearson correlation coefficients are listed in Supplementary table S1. **c.** Top 10 enriched gene sets from Gene Ontology term-based gene set enrichment analysis of correlation coefficients shown in **b**. Color shade indicates the percentage of MitoCarta3.0 genes in each gene set. **d.** Immunofluorescence analysis of N6AMT1 localization. HeLa cells were transduced with N6AMT1-3xFLAG cDNA and immunolabelled with anti-FLAG and anti-TOM20 (a mitochondrial marker). Nuclei were visualized by Hoechst staining. **e.** Immunoblot analysis of N6AMT1 protein levels in control (sgCtrl) and *N6AMT1*-depleted (sg*N6AMT1*) K562 cells. **f.** Transcriptomic analysis of 11,285 transcripts in *N6AMT1*-depleted K562 cells highlighting MitoCarta3.0 (orange) and mitochondrial DNA (mtDNA)-encoded transcripts (blue). False discovery rate (FDR) was calculated according to the method of Benjamini and Hochberg. n = 3 independent lentiviral transductions per condition. **g.** Representative basal oxygen consumption rate (OCR) and extracellular acidification rate (ECAR) per 10,000 cells in *N6AMT1*-depleted K562 cells. Bars represent mean values (n = 15 replicate wells, representative of ≥ 3 independent experiments), and error bars represent standard errors of the mean.

*Since N6AMT1* is involved in histone H4 monomethylation and possibly DNA methylation, we reasoned that it could impact mitochondria through transcriptional regulation in the nucleus. We depleted *N6AMT1* using CRISPR/Cas9 in human chronic myelogenous leukemia K562 cells and performed RNA sequencing (RNA-Seq) on total RNA (Figures 1E-F and S1C, Supplementary table S3). Gene set enrichment analysis (GSEA)^35,36^ revealed no trend in the overall transcription of MitoCarta3.0 genes (Figure S1C, Supplementary table S4) and we obtained similar results when we re-analyzed a published RNA-Seq dataset of A549 lung carcinoma cells treated with small interfering RNAs targeting *N6AMT1*^28^ (Figure S1D, Supplementary table S5). Importantly, and in contrast to earlier studies, we included genes encoded by the mitochondrial genome in our RNA-Seq analysis, allowing us to observe a profound decrease in all 13 mt-mRNAs in *N6AMT1*-depleted cells (Figure 1F). Accordingly, we found decreased respiration in *N6AMT1*-depleted cells (Figure 1G). We conclude that *N6AMT1* is required for mt-DNA expression and mitochondrial function, even though the protein is not present within mitochondria and is not directly involved in transcriptional regulation of the nuclear genes encoding mitochondrial proteins.

### *N6AMT1* depletion reduces cytosolic translation of the mt-RNA processing machinery

Given that a direct role for *N6AMT1* in nuclear transcription is unlikely to account for the changes seen in mt-DNA gene expression, we investigated the influence of *N6AMT1* on cytosolic translation using ribosome profiling (Ribo-Seq). In K562 cells, *N6AMT1* depletion led to no or marginal changes in rRNA abundance, polysome profiles, and ribosome profiles (Figures 2A-B and S2A). In contrast to earlier studies in yeast and bacteria^23,24^, we observed no appearance of halfmers in our polysome profiles (Figure 2B) and our meta-analysis of all transcripts in *N6AMT1*-depleted cells showed no difference in ribosome occupancy on termination codons (Figure 2C), indicating no global effect on polysome formation or on translation termination, respectively. However, 1,401 individual genes were differentially translated (FDR-q < 0.05) in *N6AMT1*-depleted cells (Figure 2D, Supplementary table S6). Gene ontology analysis revealed several affected pathways (Figure S2B and Supplementary table S7), and among them, we found a significant reduction in the translation of genes involved in mt-RNA metabolism, including PRORP, the catalytic subunit of the mitochondrial RNAse P, and TRMT10C, a methyltransferase also part of the mitochondrial RNAse P complex (Figure 2D). This was in marked contrast to the RNA-Seq data, which showed no difference in the steady-state levels of mt-RNA metabolism-related transcripts, including in PROPR and TRMT10C (Figure S2B, E). GSEA using the MitoPathways gene sets^32^ confirmed “mt-RNA metabolism”, “mt-RNA processing”, and “mt-RNA granules” as the three most depleted mitochondrial pathways (Supplementary table S8), a finding that we corroborated by analyzing translation efficiency (TE), a metric in which ribosome-protected mRNA fragments are normalized to transcript abundance^37^ (Supplementary tables S9-S10).

**Figure 2.**
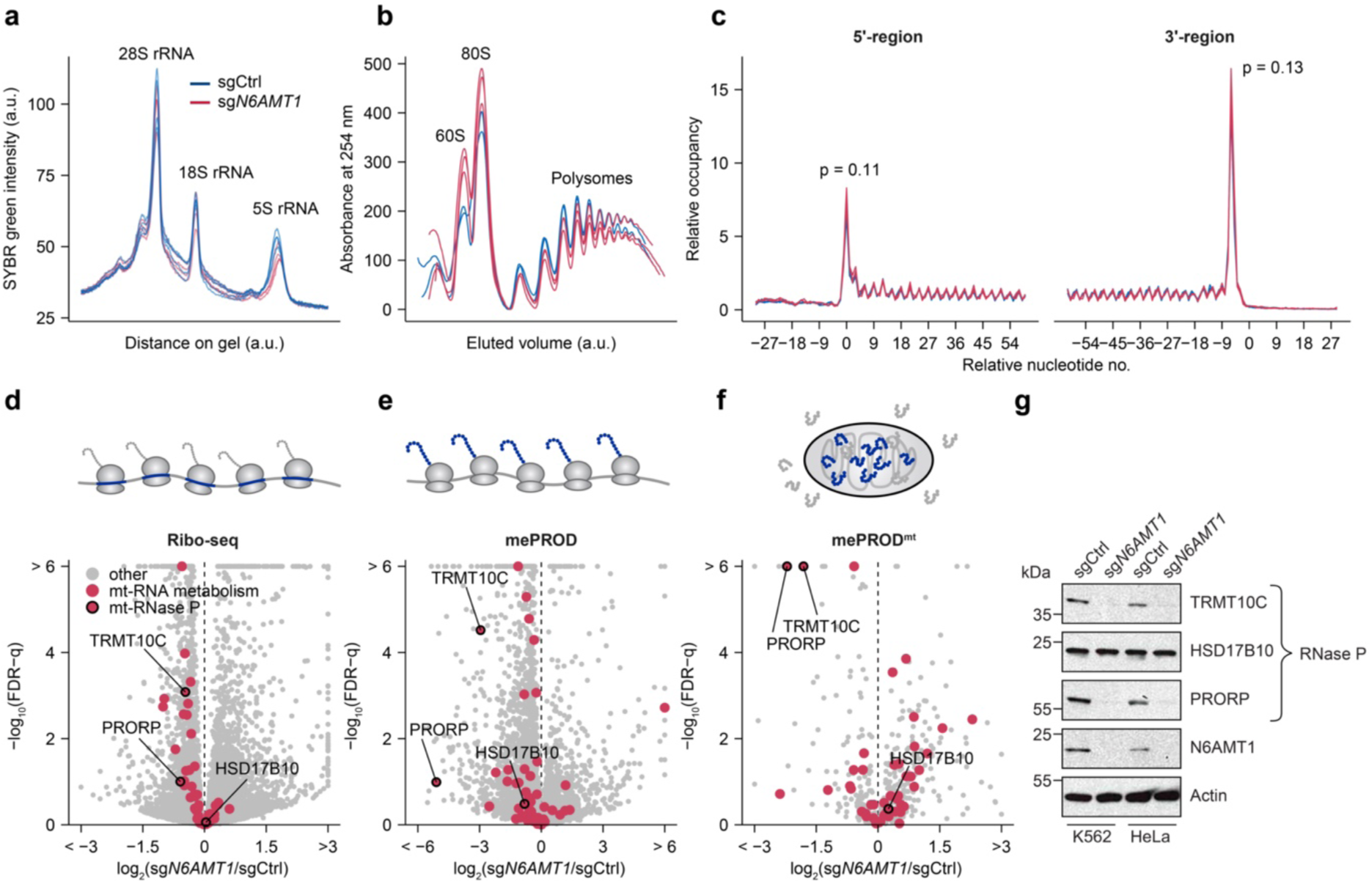
Cytosolic translation of the mt-RNA processing machinery is reduced in *N6AMT1*-depleted cells. **a.** Quantification of RNA agarose gel (shown in Figure S2A) comparing control (sgCtrl) to *N6AMT1*-depleted (sg*N6AMT1*) K562 cells 10 days post sgRNA transduction (n = 4 independent lentiviral infections per condition). Each curve represents the SYBR-green intensity across one lane, with the major ribosomal RNA (rRNA) peaks indicated. **b.** Polysome analysis of control and *N6AMT1*-depleted K562 cells (n = 2-3 lentiviral infections) 10 days post sgRNA transduction. Elution volumes are normalized to the 80S peak and the maximal polysome peak to allow direct comparisons between replicates. **c.** Metagene analysis of control and *N6AMT1*-depleted K562 cells 10 days post sgRNA transduction detected by ribosome profiling (Ribo-Seq). Curves represent ribosome-protected fragments at the indicated nucleotide positions relative to the total number of ribosome-protected fragments (n = 3 independent lentiviral infections per condition, based on 9,600 transcripts). p-values were obtained by Student’s t-tests of the 5’ and 3’ peaks, respectively. **d.** Transcript ribosome occupancy of 10,793 transcripts detected by ribosome profiling (Ribo-Seq) in *N6AMT1*-depleted K562 as compared to control cells. n = 3 independent lentiviral transduction per condition. **e-f.** Translatome analysis using multiplexed enhanced protein dynamics (mePROD; 4,079 proteins) proteomics (**e**) and global mitochondrial protein import proteomics (mePROD^mt^; 495 proteins) (**f**) in *N6AMT1*-depleted K562 as compared to control cells. n = 4 independent lentiviral transduction per condition. **g.** Protein immunoblot showing mitochondrial RNase P subunit protein levels in control and *N6AMT1*-depleted K562 and HeLa cells, with antibodies targeting TRMT10C (MRPP1), HSD17B10 (MRPP2), and PRORP (MRPP3).

We next sought to complement our ribosome profiling observations with a protein-based assay and used mePROD (multiplexed enhanced protein dynamic) and its mitochondria equivalent, mePROD^mt^ ^38,39^. mePROD combines pulsed stable isotope labeling (SILAC) with tandem mass tags, signal amplification, and mass spectrometry to monitor newly synthesized proteins on a proteome-wide scale, while mePROD^mt^ includes a mitochondrial isolation step to quantify more selectively the proteins imported into mitochondria. mePROD revealed *N6AMT1*-dependent alterations in cytosolic translation that correlated moderately with Ribo-Seq data (Pearson correlation coefficient = 0.42, Figure S2C), revealing a subset of mt-RNA-related proteins reduced in *N6AMT1*-depleted cells, including PRORP and TRMT10C (Figures 2E and S2E, Supplementary table S11). We confirmed these observations with mePROD^mt^ data (Figures 2F and S2E, Supplementary table S12). Comparing mePROD and mePROD^mt^ revealed no difference in protein import into mitochondria (Figure S2D). PRORP and TRMT10C were the two main mt-RNA-related factors depleted in all three datasets, and to confirm our findings we immunoblotted *N6AMT1*-depleted cells and observed a significant reduction in the steady-state levels of the two proteins, both in K562 and in HeLa cells, while the third RNase P subunit HSD17B10 was not affected (Figure 2G). Taken together, our complementary approaches revealed a selective reduction in cytosolic translation of two protein subunits of the mitochondrial RNAse P in *N6AMT1*-depleted cells.

### *N6AMT1* is necessary for mt-RNA processing, protein synthesis, and OXPHOS

The observed reductions in mt-DNA-encoded mRNAs (Figure 1F) and in the proteins involved in mitochondrial RNA processing (Figure 2G) prompted us to characterize further the impact of *N6AMT1* on the mitochondrial transcriptome. We used NanoString^40^ and a set of mitochondria-specific probes called “MitoString”^41^ to quantify mt-mRNAs, unprocessed mt-RNA junctions, and mt-ncRNAs (Figure S3A-B). We observed a dramatic increase in unprocessed mRNA-tRNA and rRNA-tRNA junctions in *N6AMT1*-depleted cells, affecting both 5’ and 3’ RNA junctions, as well as an accumulation of mt-ncRNAs (Figure 3A), strongly suggesting a deficiency in mt-preRNA processing, with no effect on mt-DNA copy number (Figure 3B). To exclude off-target effects, we reintroduced an sgRNA-resistant cDNA of *N6AMT1* and observed a full rescue of the phenotype (Figure 3A, C). In contrast, we found that a catalytic inactive N6AMT1 mutant (D77A^24^) failed to rescue mt-RNA processing (Figures 3C and S3C), indicating that the catalytic activity of N6AMT1 is required for its function in mt-RNA processing. To test whether PRORP and TRMT10C were limiting in the absence of *N6AMT1*, we co-expressed both cDNAs in *N6AMT1*-depleted K562 cells and observed a full or partial rescue of the mt-RNA processing phenotype at all mt-preRNA junctions tested (Figure 3D and S3D, E).

**Figure 3.**
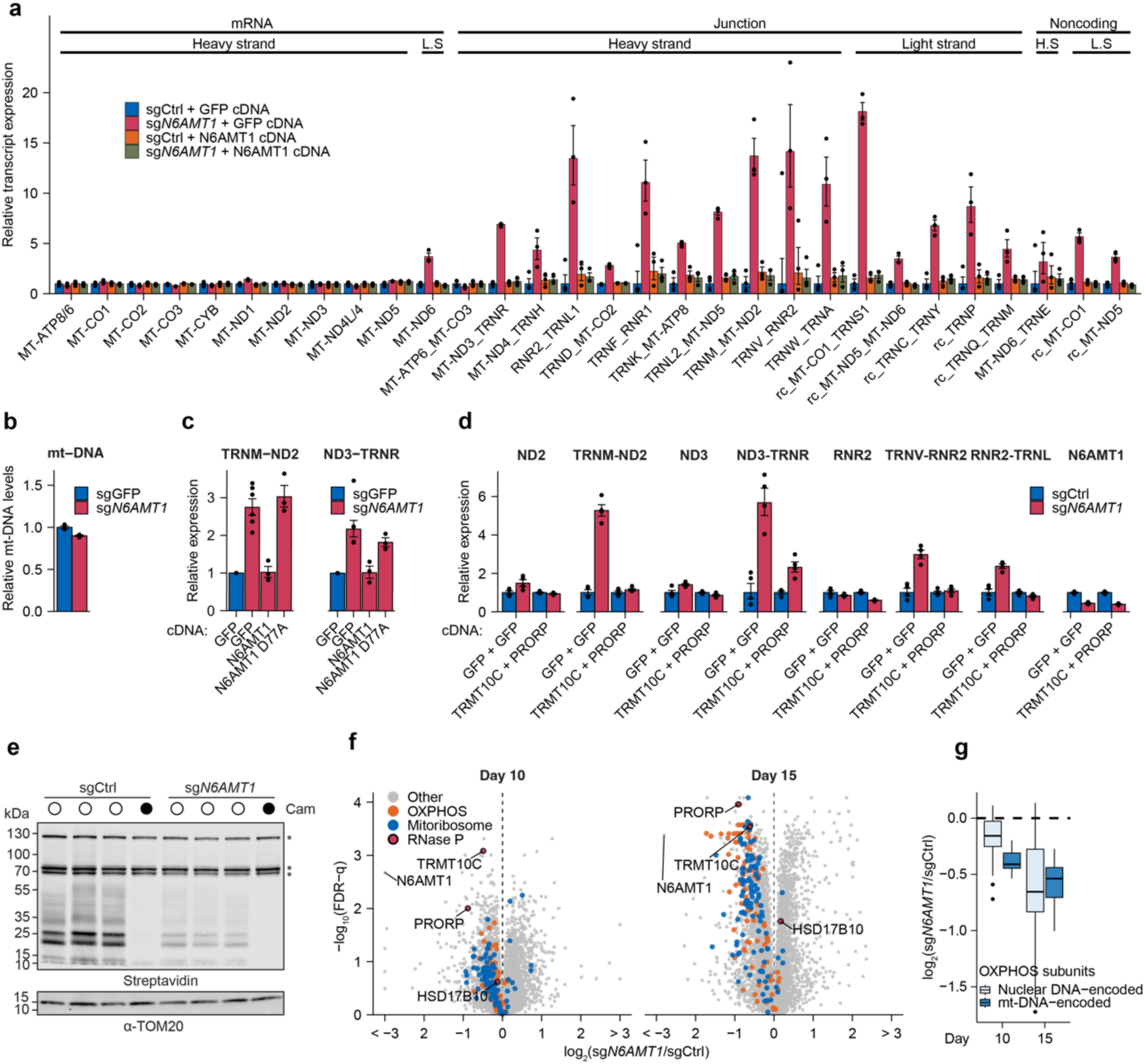
*N6AMT1* depletion leads to impaired mt-RNA processing and prevents mitochondrial translation and OXPHOS. **a.** MitoString analysis of mitochondrial RNA processing in control (sgCtrl) and *N6AMT1*-depleted (sg*N6AMT1*) K562 cells where a control cDNA (coding for GFP) or an sgRNA-resistant cDNA of *N6AMT1* were introduced. Bars represent geometric mean values and error bars indicate corresponding standard errors of the mean (n = 3 independent transduction). L.S.: light strand; H.S.: heavy strand. rc: reverse complement. MitoString probes are described in more detail in Figure S3. **b.** Relative mt-DNA quantification as determined by qPCR in control or *N6AMT1*-depleted K562 cells. Relative mt-DNA levels were calculated as the ratio between the expression of the mt-DNA-encoded MT-ND2 gene and the nuclear DNA-encoded 18S rRNA gene. **c.** qPCR analysis of unprocessed mt-RNA in control, *N6AMT1*-depleted cells, and *N6AMT1*-depleted cells with an sgRNA-resistant *N6AMT1* cDNA introduced (wild type or catalytically active mutant D77A) (n = 3-6 independent lentiviral infections). **d.** qPCR analysis of unprocessed and total mt-mRNA and mt-rRNA in control or *N6AMT1*-depleted K562 with theintroduction of either 2 different *GFP* cDNAs or *TRMT10C* + *PRORP* cDNA (n = 3 independent lentiviral infections). **e.** Translation of mitochondrially encoded proteins in control or *N6AMT1*-depleted K562 cells after labeling with L-Homopropargylglycine in the presence of cycloheximide (a cytosolic translation inhibitor). Chloramphenicol (Cam, a mitochondrial translation inhibitor) is used as control. Asterisks indicate endogenously biotinylated mitochondrial proteins. All mt-DNA-encoded proteins migrate below the 70 kDa marker. **f.** Proteome changes of 7,364 quantified proteins in *N6AMT1*-depleted K562 cells as compared to control, 10 days (left panel) or 15 days (right panel) following sgRNA transduction (n = 3 independent transduction per condition per day). **g.** Proteomics protein abundance of OXPHOS subunits according to the genome (nuclear or mitochondrial (mt-DNA) that encodes them.

MitoString was originally established and tested by knock-down of 107 mitochondrially-localized, predicted RNA-binding proteins^41^, allowing us to perform unsupervised clustering to identify which of these genes share similarities with *N6AMT1*. Among the 107 knockdowns, we found that *N6AMT1* depletion clustered best with depletion of subunits from the mitochondrial RNAse P and RNAse Z (Figure S3F), thus functionally supporting our translatome results (Figure 2D-F).

We then sought to characterize the mitochondrial consequences of low RNase P activity in *N6AMT1*-depleted cells. Consistent with defects in RNA processing and depletion of mature mt-RNA, we observed a global attenuation of translation by the mitochondrial ribosomes (Figure 3E). We confirmed this observation using time-resolved quantitative proteomics on *N6AMT1*-depleted cells, where we observed a strong, gradual depletion of most proteins involved in these processes (Figure 3F, Supplementary table S13). Our proteomics analysis also confirmed decreased levels of PRORP and TRMT10C, while HSD17B10 levels were only marginally affected (Figure 3F). For the respiratory chain, we observed a gradual reduction in all OXPHOS complexes, with the exception of complex V, which can partially assemble even in the absence of mt-DNA^42^ (Figure S3G). Consistent with the defect caused by abnormal mt-RNA expression, we noted that at early time points the mt-DNA-encoded OXPHOS subunits, which are all core subunits of the respiratory chain, appeared to be more dramatically reduced than the nuclear DNA-encoded subunits (Figure 3G). In addition, pathways downstream of mitochondrial gene expression were also progressively decreased at the protein level, as expected from their strict dependency on intact mt-RNA metabolism (Figure S3G), while mitochondrial pathways such as “signaling”, “protein import, sorting, and homeostasis”, and “metabolism” were not affected, indicating that mitochondrial functions not related to gene expression or OXPHOS were maintained (Figure S3G). We conclude that defects in *N6AMT1*-dependent synthesis of the mitochondrial RNAse P machinery alter the mitochondrial proteome in a way that leads to a progressive decline in pathways downstream of mt-RNA processing, such as mitochondrial translation and OXPHOS.

### Impaired mt-RNA processing in *N6AMT1*-depleted cells causes MRG stress and dsRNA accumulation

Mitochondrial RNA processing takes place in MRGs^5,43^, where the mitochondrial RNAses P/Z and the degradosome cleave and degrade mt-RNA. In the absence of these events, the accumulation of mt-RNA, often in the form of mt-dsRNA, can trigger an interferon response and inflammation^11^. Based on its effects on mt-preRNA and mt-ncRNA processing (Figure 3A), we hypothesized that *N6AMT1* depletion could trigger the accumulation of unprocessed mt-RNA in MRGs, as well as innate immune signaling related to mt-dsRNA. K562 cells lack a large genomic region including the interferon gene locus^44^ and, in addition, are poorly compatible with imaging. We therefore used HeLa cells to investigate the MRGs following *N6AMT1* depletion, where we also confirmed strong depletion of PRORP and TRMT10C (Figure 2G). We immunolabelled MRGs with antibodies against FASTKD2, an MRG specific marker whose abundance was not affected in our proteomics profiles (Supplementary table S13), as well as against BrU following a short pulse of 5-bromouridine (BrU) to label newly synthesized mt-RNA. We observed a striking accumulation of MRGs in *N6AMT1*-depleted cells, indicating MRG stress^5^ (Figure 4A,C). Similarly, and as expected from the accumulation of mt-ncRNA (Figure 3A), we also observed strong accumulation of dsRNA in mitochondria from *N6AMT1*-depleted cells, here too forming more abundant, larger and brighter foci following immunolabeling using a dsRNA antibody (Figure 4B,D). Finally, we examined the physiological consequences of this mt-RNA accumulation using qPCR and, as expected from previous work on patients’ cells unable to degrade mt-RNA^11^, found a transcriptional activation of innate immunity genes characterized by high expression of interferon beta (*IFNB1*) and interferon-related genes such as *USP18*, *IFIT1* or *GBP2* (Figure 4E). We conclude that the aberrant mt-RNA processing seen in *N6AMT1*-depleted cells results in the accumulation of unprocessed and double-stranded mt-RNA, with consequences for MRGs, OXPHOS, and immune signaling (Figure 4F).

**Figure 4.**
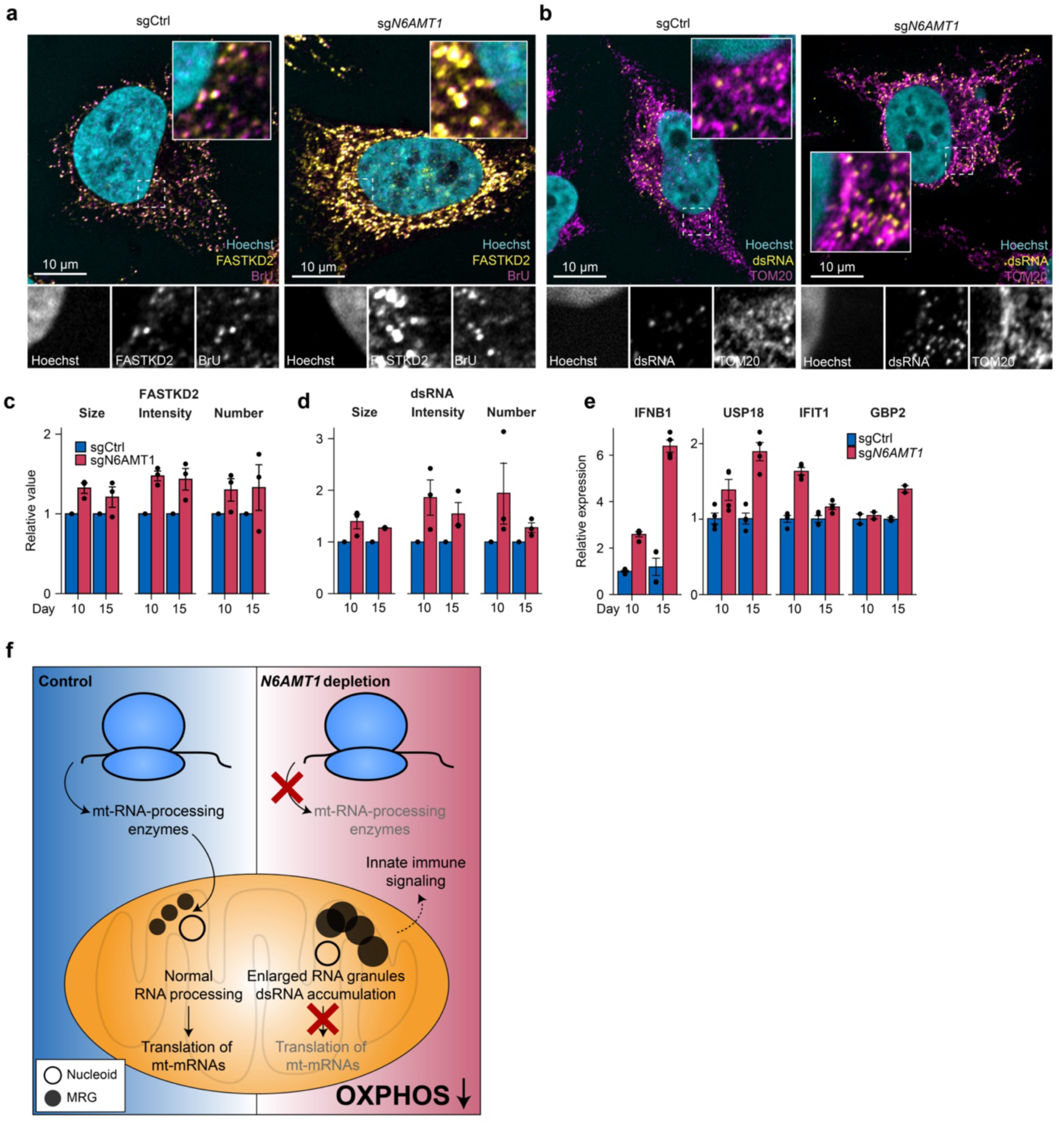
Mitochondrial preRNA and dsRNA accumulate and trigger an interferon response in *N6AMT1*-depleted cells. **a.** Representative images from immunofluorescence analysis of FASTKD2- and bromouridine (BrU)- immunolabeled mitochondrial RNA granules (MRGs) in control (sgCtrl) or *N6AMT1*-depleted (sg*N6AMT1*) HeLa cells 15 days post transduction. **b.** Representative images from immunofluorescence analysis of mitochondria (labeled with antibodies against TOM20) and mitochondrial double-stranded RNA (dsRNA) (labeled with J2 dsRNA antibody) in sgCtrl or sg*N6AMT1* HeLa cells 15 days post transduction. **c,d.** Quantification of FASTKD2 foci (c) and dsRNA foci (d), 10 or 15 days following transduction of HeLa cells with sgCtrl or sg*N6AMT1*. n = 3 independent experiments, with signal from at least 25 cells per condition. “Number” refers to average number of foci per cell area, “Size” refers to the average foci area, and “Intensity” refers to the average foci intensity. **e**. qPCR determination of transcripts of interferon beta (IFNB1) and interferon-inducible USP18, IFIT1, and GBP2 in *N6AMT1*-depleted HeLa cell lines as compared to controls 10 or 15 days post transduction. n = 4 independent transductions. **f.** Proposed role of *N6AMT1* in mitochondrial biogenesis. In *N6AMT1*-depleted cells, translation of mt-RNA-processing enzymes is reduced, leading to accumulation of unprocessed RNA in mitochondrial RNA granules. This prevents translation of essential genes involved in mitochondrial gene expression and oxidative phosphorylation (OXPHOS). In addition, impaired RNA processing leads to the accumulation of double-stranded mt-RNA that activates immune signaling pathways.

## Discussion

We report here a cytosolic pathway required for mitochondrial biogenesis that relies on N6AMT1. We show that N6AMT1 is required for the cytosolic translation of two key subunits from the mitochondrial RNA processing machinery, the deficiency of which leads to MRG stress, double-stranded RNA accumulation, immune signaling, and a collapse in OXPHOS (Figure 4F).

Our results shed light on the cellular function of N6AMT1, a multi-substrate methyltransferase with previously assigned functions ranging from transcription to arsenic metabolism. Using Ribo-Seq and mePROD we established a role for human N6AMT1 in cytosolic protein synthesis. Interestingly, N6AMT1 orthologs in bacteria and yeast are also involved in translation, and while we report that the molecular function of N6AMT1 in humans has diverged, as it does not involve ribosome assembly or translation termination (Figure 2A, B), our results nevertheless indicate that N6AMT1 has retained a canonical role in protein synthesis that appears to depend on its catalytic activity (Figure 3C). Previous work identified translation-related factors, such as RRP1, EIF2BD, and eRF1^22,25,45,46^, as substrates for N6AMT1, while in yeast, Mtq2 interacts with ribosomal proteins^24^. Further work will be necessary to clarify which of the potential N6AMT1 methylation targets accounts for its role in translation regulation.

Based on our genome-wide identification of the genes required for mitochondrial function^12^, our work reveals a pathway which is necessary for the coordinated expression of the nuclear and mitochondrial genomes. We show that *N6AMT1* is required for the translation of PRORP and TRMT10C, two subunits of the mitochondrial RNAse P which are central for mitochondrial gene expression, and which are also implicated in mitochondrial disease^47,48^. Here we have focused our investigations on these two mitochondrial RNase P subunits and report that their decreased translation impairs mitochondrial RNA processing with two major consequences: (1) mt-mRNAs, mt-rRNAs and mt-tRNAs remain unprocessed, thus prohibiting mitochondrial ribosome assembly and translation, strongly reducing expression of the 13 mt-DNA-encoded proteins and inhibiting OXPHOS (Figure 2); (2) mt-RNA accumulates in mitochondria, where it colocalizes with markers of MRGs. Based on genetic depletion, our work thus far points to a role for *N6AMT1* in supporting fully functional mitochondria, and it is possible that N6AMT1 may be induced, or activated, during intense mitochondrial biogenesis programs, such as those seen during adipose or muscle differentiation. In the future, it will be interesting to analyze changes in *N6AMT1* expression during such processes, and to follow the expression of the mitochondrial RNAse P, mt-RNA processing, and the state of MRGs.

Recent work has highlighted the role of both mt-DNA and mt-RNA in triggering inflammation and the interferon response^11,49,50^. In the case of mt-RNA, depletion or mutations in *PNPT1* lead to mt-dsRNA accumulation in mitochondria and its subsequent leakage into the cytosol where it serves as ligand for MDA-5 and RIG-I, leading to both NF-κB and interferon signaling^11^. While our K562 cellular model used here for large-scale genetic experiments did not allow us to study interferon signaling, we detected significant up-regulation of IFNB1 mRNA as well as interferon-related transcripts in *N6AMT1*-depleted HeLa cells (Figure 4E), a result very similar to that observed following *PNPT1* depletion or mutation^11^.

High levels of N6AMT1 are associated with multiple forms of cancers, and small interfering RNAs to *N6AMT1* prevent the proliferation of cancer cells from androgen receptor–dependent and castration-resistant prostate cancer^27^, bladder cancer^51,52^, colorectal cancers^53^, as well as small cell and non-small cell lung cancer^28^. N6AMT1 methyltransferase inhibitors are also being developed^54,55^, and early results show promising activity at blocking tumor cell proliferation. A proposed mechanism for the role of N6AMT1/KMT9 in cancer is the monomethylation of histone H4 that controls transcription of cell cycle regulators and pro-proliferation factors^27^. We have reported here on the role of *N6AMT1* in cytosolic translation, mitochondria, and innate immunity. Mitochondria are central to the provision of energy and building blocks that support cell proliferation, including in cancer cells. Based on our observations, we propose that the mechanism by which *N6AMT1* promotes cancer cell growth also involves mitochondrial activity. This role is supported by our analysis of *N6AMT1* in 1,100 CRISPR/Cas9 screens performed across cancer cell lines from 20 tissues of origin, where we report that the *N6AMT1* dependency profile correlates best with dependency on mitochondria (Figure 1A-C). Our observations are compatible with earlier studies reporting a function of N6AMT1/KMT9 in epigenetics^27,53^, and, collectively, the evidence points to a dual role for N6AMT1 in both gene expression and protein synthesis. Going forward, it will be important to identify which of these functions underlie cancer cell proliferation, as this could promote the development of novel anti-neoplastic molecules with reduced drug toxicity.

## Acknowledgements

The authors would like to thank past and present members of the Jourdain lab as well as S. Chamois, D. Gatfield, R. Green, V. Heurgué-Hamard, A. Jeltsch, J.C. Martinou, E. Mick, D. Mohr, H.P. Nadimpalli, C.C. Wu, G.L. Xu, the UNIL Protein Analysis Facility, the members of the Center for Function Proteomics at Goethe University Frankfurt, and the Johns Hopkins Genetic Resources Core Facility. This work was supported by grants from the Swiss National Science Foundation (310030_200796 to AAJ) and the Hessian Ministry for Arts and Sciences excellence initiative EnABLE (to CM). VKM is an Investigator of HHMI. BZ was an HHMI fellow of the Damon Runyon Cancer Research Foundation (DRG-2250-16) during the course of this study.

## Material and Methods

### Cell lines

K562 (CCL-243), HeLa (CCL-2) and 293T (CRL-3216) were obtained from ATCC and re-authenticated by STR profiling at ATCC during the course of this study.

### Cell culture

Cells were maintained in DMEM containing 1 mM sodium pyruvate (ThermoFisher Scientific) with 25 mM glucose, 10% fetal bovine serum (FBS, ThermoFisher Scientific), and 100 U/ml penicillin/streptomycin (ThermoFisher Scientific) under 5% CO_2_ at 37°C. Cells were counted using a ViCell Counter (Beckman) and only viable cells were considered.

### Cancer Cell Line correlation analysis

DepMap Public 23Q2 Chronos scores were obtained from the Cancer Cell Line Encyclopedia (CCLE) Dependency Portal (DepMap)^56,57^. Genes annotated as common essentials or with missing values were excluded. Pearson’s correlation coefficients were calculated between N6AMT1 Chronos scores and all other genes using R version 4.3.1^58^. Gene-wise correlation coefficients were exported for GSEA.

### Analysis of published RNA-Seq datasets

Count values from dataset GSE131016 were obtained from the NCBI Gene Expression Omnibus data repository and re-analyzed using DESeq2^59^ in R version 4.3.1^58^. For GSEA, log_2_ fold change shrinkage was perfomed using the apeglm package version 1.26.1^60^.

### Gene set enrichment analysis

CCLE dependency correlation coefficients and log_2_(fold-changes) from RNA-Seq, Ribo-Seq, and proteomics analyses were analyzed by GSEA version 4.3.2^36^ using the c5.go.v2023.Hs.symbols (Gene ontology) or MitoPathways^32^ gene sets.

### Gene-specific CRISPR/Cas9 knockouts and cDNA rescue

The two best *N6AMT1* sgRNAs from the Avana-library^61^ were ordered as complementary oligonucleotides (Integrated DNA Technologies) and cloned in pLentiCRISPRv2^62^. sgRNAs targeting EGFP (non-targeting) or *OR11A1* and *OR2M4* (two unexpressed genes in K562 cells) were used as a negative control. Lentiviruses were produced according to Addgene’s protocol and 24-48 h post-infection cells were selected with 2 µg/ml puromycin (ThermoFisher Scientific) for 2-5 days. Cells were then maintained in routine culture media for 10-15 days post transduction before analysis. Gene disruption efficiency was verified by immunoblotting. For rescue, a sgRNA-resistant version of N6AMT1-3xFLAG cDNA was synthesized in pWPI (Addgene). The same cDNA was further edited to obtain the D77A variant using site-directed mutagenesis. sgRNA, cDNA and primer sequences are described in Supplementary table S14.

### Mitochondrial and nuclear DNA determination

Mitochondrial and nuclear DNA determination was carried as previously described^32^. Approximately 1x10^5^ cells from each condition were harvested and lysed in 100 μl mt-DNA lysis buffer (25mM NaOH, 0.2 mM EDTA) before incubation at 95 °C for 15min. 100 μl of 40 mM Tris-HCl pH 7.5 was added to neutralize the reaction on ice. Samples were diluted 50x and the ratio between mitochondrial and nuclear DNA was determined with a CFx96 quantitative PCR machine (Bio-rad) using probes and primers targeting MT-ND2 and 18S rRNA genes as described in Supplementary table S15. Relative mt-DNA abundance was determine using the ΔΔCt method.

### RNA extraction, reverse transcription, and quantitative PCR (qPCR)

RNA was extracted from total cells with the RNeasy kit (QIAGEN) and digested by DNase I before reverse transcription with murine leukemia virus reverse transcriptase using random primers (Promega). qPCR was performed with and a CFx96 Touch Real-Time PCR system (Bio-rad) or a LightCycler 480 II (Roche) using probes and primers described in Supplementary table S15. All data were normalized to the expression of TATA-box binding Protein (TBP) using the ΔΔCt method.

### Seahorse

1.25x10^5^ K562 cells were plated on a Seahorse plate in Seahorse XF DMEM media (Agilent) containing 25 mM glucose and 4 mM glutamine (ThermoFisher Scientific). Oxygen consumption and extracellular acidification rates were simultaneously recorded by a Seahorse XFe96 Analyzer (Agilent) using the Mito Stress Test protocol, in which cells were sequentially perturbed by 2 µM oligomycin, 1 µM CCCP, and 0.5 µM antimycin (Sigma). Data were analyzed using the Seahorse Wave Desktop Software (Agilent). Data were not corrected for carbonic acid derived from respiratory CO_2_.

### Immunofluorescence and confocal microscopy

For immunofluorescence, cells transduced with N6AMT1-3xFLAG, sgCtrl or sgN6AMT1 sgRNAs were successively fixed in 4% paraformaldehyde in cell culture media at room temperature for 10 min, permeabilized for 15 min in 0.2% Triton-PBS solution (Sigma-Aldrich, T8787), incubated with primary antibodies (1:200) in 1% PBS-Bovine Serum Albumin (Sigma Aldrich, A9647-100G) for 1 h, washed 3x5 min in 1% Bovine Serum Albumin (BSA) in PBS, incubated with fluorophore-coupled secondary antibodies (1:300) and Hoechst (1:1000) in 1% BSA in PBS for 45 min, washed 3x in PBS and mounted on a slide using FluorSave (EMD Millipore). Cells were imaged using Zeiss LSM 880 Airyscan confocal microscope (Zeiss, 518F). For bromouridine (BrU) staining HeLa cells were pulsed with 5 mM BrU for 30 minutes before fixation. MRG and dsRNA quantification was performed using an automated pipeline on CellProfiler software. To limit our quantification to the mitochondrial network, nuclei were removed from the analysis and cell delimitation was performed using a mitochondrial marker.

### Polyacrylamide gel electrophoresis and immunoblotting

Cells were harvested, washed in PBS and lysed for 5 min on ice in RIPA buffer (25 mM Tris pH 7.5, 150 mM NaCl, 0.1% SDS, 0.1% sodium deoxycholate, 1% NP40 analog, 1X protease inhibitor (Cell Signaling) and 1:500 Universal Nuclease (ThermoFisher Scientific)). Protein concentration was determined from total cell lysates using DC Protein Assay (Bio-rad). Gel electrophoresis was done on Novex Tris-Glycine gels (ThermoFisher Scientific) before transfer using the Trans-Blot Turbo blotting system and nitrocellulose membranes (Bio-rad). All immunoblotting was performed in 5% milk powder or 5% BSA in Tris-Buffered Saline (TBS; 20 mM Tris, pH 7.4, 150 mM NaCl)+ 0.1% Tween-20 (Sigma). Washes were done in TBS + 0.1% Tween-20 (Sigma). Specific primary antibodies were diluted 1:1000-1:5000 in blocking buffer (TBS + 0.1% Tween-20 and 5% milk powder). Fluorescent- and HRP-coupled secondary antibodies were diluted 1:10,000 in blocking buffer. Membranes were imaged with an Odyssey CLx analyzer (Li-Cor) or chemiluminescence and films.

### Antibodies

Antibodies were: N6AMT1 (LSBio LS-C346278), actin (Sigma A1978), FASTKD2 (Proteintech 17464-1-AP), TRMT10C (Sigma, HPA036671), HSD17B10 (Abcam, ab10260), PRORP (Abcam, ab185942), GAPDH (CST, 5174T), bromouridine (Sigma 11170376001), dsRNA (Exalpha, 10010500), TOM20 (CST, 42406).

### L-Homopropargylglycine labeling

15 days post transduction with control sgRNA or *N6AMT1*-targeted sgRNA, K562 cells were washed with prewarmed PBS and resuspended in methionine-free medium (DMEM, high glucose, no glutamine, no methionine, no cystine (ThermoFisher Scientific) with 200 µM L-cystine, 4 mM glutamine, and 1 mM sodium pyruvate). Mitochondrial and cytosolic translation was inhibited by incubation with 100 µg/ml chloramphenicol (Sigma-Aldrich) and 50 µg/ml cycloheximide (Sigma-Aldrich), respectively, 10 min before the addition of 500 µM L-homopropargylglycine (Jena Bioscience). After 1.5 h, cells were harvested, washed once in cold PBS then twice in cold mitochondrial isolation buffer (10 mM HEPES pH 7.4, 210 mM mannitol, 70 mM sucrose), and mitochondria were isolated as previously described^63^. In short, 15x10^6^ cells were resuspended in 750 µl MB + 250 U/ml Pierce Universal Nuclease (Thermo Fisher Scientific) and 1X protease inhibitor (Cell Signaling) and lysed with 10 strokes of a 25 G needle. Lysates were centrifuged 10 min at 500 *g*, 4 °C to remove debris and intact cells, and supernatants were further centrifuged 10 min at 10,000 *g*, 4 °C to pellet mitochondria. Mitochondrial pellets were dissolved in 60 µl RIPA buffer, and protein concentration was determined using a DC Protein Assay (Bio-Rad). 50 µl dissolved mitochondrial pellets were mixed with 50 µl click-reaction mix (10 mM sodium ascorbate (Jena Bioscience); 40 µM picolyl-Azide-PEG4-Biotin (Jena Bioscience); 2.4 mM 2-(4-((bis((1-(tert-butyl)-1H-1,2,3-triazol-4-yl)methyl)amino)methyl)-1H-1,2,3-triazol-1-yl)acetic acid (Jena Bioscience); 1.2 mM CuSO4 (Jena Bioscience)), incubated 60 min at room temperature, and mixed with 20 µl 5X SDS-PAGE loading buffer (250 mM Tris pH 6.8, 4% sodium dodecyl sulfate, 0.06% (w/v) bromophenol blue, 16% (v/v) beta-mercaptoethanol, 30% (v/v) glycerol). Biotinylated proteins were visualized by immunoblotting using IRDye 800CW Streptavidin (LI-COR).

### Total ribosomal RNA quantification by gel electrophoresis

K562 cells were harvested 10 days post transduction with control sgRNA or *N6AMT1*-targeted sgRNA, washed in cold PBS, and snap-frozen in liquid nitrogen. RNA was extracted using TRI reagent (Sigma-Aldrich), essentially according to the manufacturer’s instructions, with the inclusion of GlycoBlue Coprecipitant (Invitrogen) for improving the visibility of precipitated RNA. 16 µg purified RNA was mixed with 6X gel loading dye (New England BioLabs) and run on a 1% agarose-TAE (20 mM Tris; 10 mM acetic acid; 0.5 mM EDTA) gel + 0.01% (v/v) SYBR safe DNA Gel Stain (Invitrogen). SYBR green signal intensity was quantified across each lane using ImageJ version 1.53.

### Polysome profiling

10 days post viral transduction, control or sgN6AMT1-infected K562 cells were treated with 100 µg/ml cycloheximide 10 min before harvesting. Cells were harvested, washed in PBS, and lysed for 10 min on ice in lysis buffer (50 mM HEPES pH 7.3, 100 mM potassium acetate, 15 mM magnesium acetate, 5% glycerol, 0.5% Triton X-100, 2X protease inhibitor (Cell Signaling), 5 mM DTT, 100 µg/ml cycloheximide). Lysates were centrifuged 9 min at 9,000 *g*, 4 °C, and RNA concentrations were measured with a Nanodrop ND-2000. 180 µg RNA was loaded on 10-50% sucrose gradients in 25 mM HEPES, 100 mM potassium acetate, 5 mM magnesium acetate and 100 µg/ml cycloheximide and centrifuged 1:45 h at 40,000 *g*, 4 °C on a Beckman Coulter Optima LE-80K Ultracentrifuge. Gradients were analyzed on an Äkta purifier measuring absorbance at 254 nm. Elution volumes were normalized to volumes at the maximal absorbance (80S peak) and the fourth polysome peak to allow direct comparison between samples.

### Next-generation RNA sequencing and ribosome profiling

3 replicates each of WT and N6AMT1 KO were performed for both RNA-Seq and ribosome profiling. For each replicate, 5x10^6^ cells were harvested by centrifugation and flash-frozen in liquid nitrogen. Frozen cell pellets were directly resuspended by pipetting in 200 μl of mammalian footprint lysis buffer (20 mM Tris-Cl pH7.4, 150 mM KCl, 5 mM MgCl_2_, 1 mM DTT, 1% Triton X-100, 2 units/ml Turbo DNase (Thermo Fisher Scientific), 0.1 mg/ml cycloheximide (Sigma-Aldrich)). Lysates were incubated on ice for 10 min and then cleared by centrifugation at 20,000 *g* for 10 min at 4 °C. Cleared lysates were quantified by a Nanodrop ND-1000 Spectrophotometer (Thermo Fisher Scientific), and aliquots of 1.5 and 0.5 A260 units were flash frozen in liquid nitrogen. For RNA-Seq, 200 µl TRIzol (Thermo Fisher Scientific) was added to an 0.5 A260 aliquot, and RNA was purified with the Zymo Direct-zol RNA miniprep kit according to the manufacturer’s instructions for purifying total RNA. Resulting RNA was quantified by a NanoDrop ND-1000 Spectrophotometer (Thermo Fisher Scientific), and RNA-Seq libraries were prepared using the Illumina TruSeq stranded total RNA-Seq kit with ribozero gold human/mouse/rat rRNA depletion, following the manufacturer’s instructions. After final PCR, DNA was quantified by Bioanalyzer High-sensitivity DNA Chip (Agilent) and equimolarly pooled. For ribosome profiling, 1.5 A260 units of lysate was diluted to 200 µl with lysis buffer and digested with 750 units of RNase I (Ambion) in 200 µl lysis buffer at 25 °C for 1 h with 500 rpm agitation in a thermomixer. Reactions were quenched with 200 U SUPERase-in (Thermo Fisher Scientific) and pelleted through a 900 µl sucrose cushion in a TLA100.3 rotor for 1 h at 100,000 rpm. Pellets were resuspended in 300 µl TRIzol (Thermo Fisher Scientific) and RNA extracted with the Zymo Direct-zol RNA miniprep kit according to the manufacturer’s instructions for purifying total RNA including small RNAs. Fragments were size selected on a 15% TBE-urea PAGE gel (Bio-Rad) with 15 and 36 nt oligos as size markers, cutting a range from 15 to 36 nt. After gel elution, fragments were dephosphorylated with PNK, and, ligated to preadenylated 3′ linker oBZ407. rRNA depletion was performed with the Ribo-Zero Gold kit (Illumina). Reverse transcription was performed using protoscript II (NEB) and RT primer oBZ408, and RNA degraded by base hydrolysis. cDNA was gel purified and circularized with Circ Ligase (Lucigen). Libraries were PCR amplified using forward primer oBZ287 (NINI2) and reverse primers bar 11 through 16 (oBZ207-212). PCR was performed with Phusion DNA polymerase, 1 µM of each primer, and 8 cycles. Amplified libraries were purified away from primers on an 8% native TBE acrylamide gel. After extraction, DNA was quantified by Bioanalyzer High-Sensitivity DNA Chip (Agilent) then equimolarly pooled. All libraries were sequenced on an Illumina HiSeq 2500 at the Johns Hopkins genetic resources core facility with 60 nt single-end reads. RNA-Seq and Ribo-Seq adapters are described in Supplementary tables S16-S17.

### RNA-Seq mapping

Fastq files were adaptor stripped using cutadapt with a minimum length of 15 and a quality cutoff of 2 (parameters: -a NNNNNNCACTCGGGCACCAAGGAC –minimum-length = 15 –quality-cutoff = 2). Resulting reads were mapped, using default parameters, with HISAT2^64^ using a GRCh38, release 84 genome and index. Differential expression analysis was performed using DESeq2^59^ with counts generated by overlapping with a GRCh38, release 84 gtf. Volcano plots were generated by plotting the resulting log_2_ fold change against a -log_10_ transformation of the FDR-adjusted (Benjamini-Hochberg) p-value.

### Ribo-Seq mapping and analysis

Fastq files were adaptor stripped using cutadapt. Only trimmed reads were retained, with a minimum length of 15 and a quality cutoff of 2 (parameters: -a CTGTAGGCACCATCAAT – trimmed-only – minimum-length = 15 –quality-cutoff = 2). Histograms were produced of ribosome footprint lengths and reads were retained if the trimmed size was 26 or 34 nucleotides. Resulting reads were mapped, using default parameters, with HISAT2^64^ using a GRCh38, release 84 genome and index and were removed if they mapped to rRNA or tRNA according to GRCh38 RepeatMasker definitions from UCSC. A full set of transcripts and CDS sequences for Ensembl release 84 was then established. Only canonical transcripts (defined by knownCanonical table, downloaded from UCSC) were retained with their corresponding CDS. Retained reads were then mapped to the canonical transcriptome with bowtie2 using default parameters. For analysis, the P-site position of each read was predicted by riboWaltz^65^ and confirmed by inspection. Counts were made by aggregating P-sites overlapping with the CDS and P-sites Per Kilobase Million (PPKMs) were then generated through normalizing by CDS length and total counts for the sample. Differential expression and translational efficiency (ribosome occupancy / mRNA abundance) analysis coupled with RNA-Seq data was performed using DESeq2^59^ with counts generated by overlapping with a GRCh38, release 84 gtf. Volcano plots were generated by plotting the resulting log_2_ fold change against a -log_10_ transformation of the adjusted p-value. All metagenes were calculated on a subset of expressed canonical transcripts which had PPKM values greater than 1 across all samples (9,600 transcripts). Within these, P-site depths per nucleotide were normalized to the mean value in their respective CDS. For metagenes around start and stop, the mean of these normalized values was taken for each nucleotide within 30 nt upstream and 60 nt downstream of start or 60 nt upstream and 30 nt downstream of stop.

### mePROD and mePROD^mt^

K562 cells were infected with sgCtrl or sg*N6AMT1* in quadruplicates (two sgRNAs per condition). 10-11 days post transduction, cells were resuspended in heavy labelled amino acids (SILAC)-containing medium for 3 h. Cells were collected and washed twice with cold PBS. For mePROD^mt^, crude mitochondrial were obtained as described previously^63^. Briefly, pellets were washed once in cold mitochondrial isolation buffer (10 mM HEPES, pH 7.4, 210 mM mannitol, 70 mM sucrose, 1 mM EDTA), and resuspended in 500 µl cold MIB. Cells were lysed by 30 strokes with a 25 G needle and diluted with 1 ml cold MIB. Lysates were centrifuged at 2,000 *g*, 5 min at 4 °C, and resulting pellets were lysed a second time as described above. Supernatants were pooled, centrifuged at 13,000 *g*, 10 min at 4 °C, and crude mitochondrial pellets were used in downstream analyses. Lysis, sample preparation and tandem-mass tag (TMT)-labelling was carried out as described previously^38^. Equal amounts of TMT-labeled peptides were pooled and fractionated into 8 fractions using a High pH Reversed-Phase Peptide Fractionation Kit (ThermoFisher Scientific) according to the manufacturer’s instructions.

For mass spectrometry analysis, Samples were analyzed with settings described earlier^39^. Briefly, fractions were resuspended in 2% acetonitrile and 0.1% formic acid and separated on an Easy nLC 1200 (ThermoFisher Scientific) and a 35 cm long, 75 μm ID fused-silica column, which had been packed in house with 1.9 μm C18 particles (ReproSil-Pur, Dr. Maisch) and kept at 50 °C using an integrated column oven (Sonation). HPLC solvents consisted of 0.1% formic acid in water (Buffer A) and 0.1% formic acid, 80% acetonitrile in water (Buffer B). Assuming equal amounts in each fraction, 1 µg of peptides were eluted by a non-linear gradient from 3 to 60% B over 125 min followed by a step-wise increase to 95% acetonitrile in 1 min which was held for another 9 min. After that, peptides directly sprayed into an Orbitrap Fusion Lumos mass spectrometer equipped with a nanoFlex ion source (ThermoFisher Scientific). Full scan MS spectra (350-1400 m/z) were acquired with a resolution of 120,000 at m/z 100, maximum injection time of 100 ms and AGC target value of 4x10^5^. For targeted mass difference-based runs, the 10 most intense ions with a charge state of 2-5 were selected together with their labeled counterparts (Targeted Mass Difference Filter, Arg and lysine delta mass, 1%-100% partner intensity range with 5 ppm mass difference tolerance), resulting in 20 dependent scans (Top20). Precursors were isolated in the quadrupole with an isolation window of 0.7 Th. MS2 scans were performed in the quadrupole using a maximum injection time of 86 ms, AGC target value of 1x10^5^. Ions were then fragmented using HCD with a normalized collision energy of 38% and analyzed in the Orbitrap with a resolution of 50,000 at m/z 200. Repeated sequencing of already acquired precursors was limited by a dynamic exclusion of 60 s and 7 ppm, and advanced peak determination was deactivated.

Raw data was analyzed with Proteome Discoverer 2.4 (ThermoFisher Scientific). SequenceHT node was selected for database searches of MS2-spectra. Human trypsin-digested proteome (Homo sapiens SwissProt database (TaxID:9606, version 2020-03-12)) was used for protein identifications. Contaminants (MaxQuant “contamination.fasta”) were determined for quality control. Fixed modifications were: TMT6 (+229.163) at the N-terminus, TMT6+K8 (K, +237.177), Arg10 (R, +10.008), and carbamidomethyl (C, +57.021) at cysteine residues. Methionine oxidation (M, +15.995) and acetylation (+42.011) at the protein N-terminus were set for dynamic modifications. Precursor mass tolerance was set to 10 ppm and fragment mass tolerance was set to 0.02 Da. Default Percolator settings in Proteome Discoverer were used to filter peptides spectrum matches (PSMs). Reporter ion quantification was achieved with default settings in consensus workflow. PSMs were exported for further analysis using the DynaTMT package^39^. Normalized abundances from DynaTMT were used for statistical analysis after removing common contaminants and peptides with only missing values. Differential expression analysis was performed using peptide-based linear mixed models (PBLMM)^39,66^.

### Quantitative steady-state proteomics

For sample preparation, samples were digested following a modified version of the iST method^67^. Based on tryptophane fluorescence quantification^68^, 100 µg of proteins at 2 µg/µl in miST lysis buffer (1% Sodium deoxycholate, 100 mM Tris pH 8.6, 10 mM DTT), were transferred to new tubes. Samples were heated 5 min at 95 °C, diluted 1:1 (v:v) with water containing 4 mM MgCl_2_ and benzonase (Merck #70746, 250 U/µl), and incubated for 15 min at RT to digest nucleic acids. Reduced disulfides were alkylated by adding 0.25 sample volumes of 160 mM chloroacetamide (32 mM final) and incubating for 45 min at RT in the dark. Samples were adjusted to 3 mM EDTA and digested with 0.5 µg Trypsin/LysC mix (Promega #V5073) for 1h at 37 °C, followed by a second 1 h digestion with an additional 0.5 µg of proteases. To remove sodium deoxycholate, two sample volumes of isopropanol containing 1% TFA were added to the digests, and the samples were desalted on a strong cation exchange (SCX) plate (Oasis MCX; Waters Corp., Milford, MA) by centrifugation. After washing with isopropanol/1% TFA, peptides were eluted in 200 µl of 80% MeCN, 19% water, 1% (v/v) ammonia, and dried by centrifugal evaporation. For peptide fractionation, aliquots (1/8) of samples were pooled and separated into 6 fractions by off-line basic reversed-phase (bRP) using the Pierce High pH Reversed-Phase Peptide Fractionation Kit (Thermo Fisher Scientific). The fractions were collected in 7.5, 10, 12.5, 15, 17.5 and 50% acetonitrile in 0.1% triethylamine (∼pH 10). Dried bRP fractions were redissolved in 50 µl 2% acetonitrile with 0.5% TFA, and 5 µl were injected for LC-MS/MS analyses. Liquid Chromatography-Mass Spectrometry analysis (LC/MS-MS) were carried out on a TIMS-TOF Pro (Bruker, Bremen, Germany) mass spectrometer interfaced through a nanospray ion source (“captive spray”) to an Ultimate 3000 RSLCnano HPLC system (Dionex). Peptides (200 ng) were separated on a reversed-phase custom packed 45 cm C18 column (75 μm ID, 100 Å, Reprosil Pur 1.9 µm particles, Dr. Maisch, Germany) at a flow rate of 0.250 µl/min with a 2-27% acetonitrile gradient in 93 min followed by a ramp to 45% in 15 min and to 90% in 5 min (total method time: 140 min, all solvents contained 0.1% formic acid). Identical LC gradients were used for data-dependent acquisitions (DDA) and data-independent acquisitions (DIA) measurements. For creation of the spectral library, DDA were carried out on the 6 bRP fractions sample pool using a standard TIMS PASEF method^69^ with ion accumulation for 100 ms for each survey MS1 scan and the TIMS-coupled MS2 scans. Duty cycle was kept at 100%. Up to 10 precursors were targeted per TIMS scan. Precursor isolation was done with a 2 Th or 3 Th windows below or above *m/z* 800, respectively. The minimum threshold intensity for precursor selection was 2500. If the inclusion list allowed it, precursors were targeted more than one time to reach a minimum target total intensity of 20,000. Collision energy was ramped linearly based uniquely on the 1/k0 values from 20 (at 1/k0 = 0.6) to 59 eV (at 1/k0 = 1.6). Total duration of a scan cycle including one survey and 10 MS2 TIMS scans was 1.16 s. Precursors could be targeted again in subsequent cycles if their signal increased by a factor 4.0 or more. After selection in one cycle, precursors were excluded from further selection for 60s. Mass resolution in all MS measurements was approximately 35,000. The DIA method used mostly the same instrument parameters as the DDA method and was as reported previously^69^.

Per cycle, the mass range 400-1200 m/z was covered by a total of 32 windows, each 25 Th wide and a 1/k0 range of 0.3. Collision energy and resolution settings were the same as in the DDA method. Two windows were acquired per TIMS scan (100 ms) so that the total cycle time was 1.7 s.

Raw Bruker MS data were processed directly with Spectronaut 17.6 (Biognosys, Schlieren, Switzerland). A library was constructed from the DDA bRP fraction data by searching the annotated human proteome (www.uniprot.org) database of 2022 (uniprot_sprot_2022-01-07_HUMAN.fasta). For identification, peptides of 7-52 AA length were considered, cleaved with trypsin/P specificity and a maximum of 2 missed cleavages. Carbamidomethylation of cysteine (fixed), methionine oxidation and N-terminal protein acetylation (variable) were the modifications applied. Mass calibration was dynamic and based on a first database search. The Pulsar engine was used for peptide identification. Protein inference was performed with the IDPicker algorithm. Spectra, peptide and protein identifications were all filtered at 1% FDR against a decoy database. Specific filtering for library construction removed fragments corresponding to less than 3 AA and fragments outside the 300-1800 m/z range. Also, only fragments with a minimum base peak intensity of 5% were kept. Precursors with less than 3 fragments were also eliminated and only the best 6 fragments were kept per precursor. No filtering was done on the basis of charge state and a maximum of 2 missed cleavages was allowed. Shared (non proteotypic) peptides were kept. The library created contained 127,456 precursors mapping to 92,961 stripped sequences, of which 88,718 were proteotypic. These corresponded to 7,860 protein groups (7,949 proteins). Of these, 790 were single hits (one peptide precursor). In total 752,440 fragments were used for quantitation. Peptide-centric analysis of DIA data was done with Spectronaut 17.6 using the library described above. Single hits proteins (defined as matched by one stripped sequence only) were kept in the analysis. Peptide quantitation was based on XIC area, for which a minimum of 1 and a maximum of 3 (the 3 best) precursors were considered for each peptide, from which the median value was selected. Quantities for protein groups were derived from inter-run peptide ratios based on MaxLFQ algorithm^70^. Global normalization of runs/samples was done based on the median of peptides. The raw output table contained 7,675 quantified protein groups (minimum 1 precursor per group) with an overall data completeness of 94.9%. All subsequent analyses were done with the Perseus software package (version 1.6.15.0)^70^. Contaminant proteins were removed, and quantited values were log_2_-transformed. After assignment to groups, only proteins quantified in at least 4 samples of one group were kept (7,372 protein groups). After missing values imputation (based on normal distribution using Perseus default parameters), t-tests were carried out between N6AMT1-depleted and control samples, with Benjamini-Hochberg correction for multiple testing.

### NanoString

A mitochondrial-specific version of NanoString called MitoString was performed as previously described^41^. All counts were normalized to TUBB. For the unsupervised clustering, mean expression values from the previously published dataset^41^ and the MitoString analysis were combined and hierarchically clustered and visualized using the pheatmap package^71^ in R with default clustering settings.

## Data availability

Steady state proteomics and mePROD proteomics data have been deposited to the ProteomeXchange consortium (www.proteomexchange.org) via the PRIDE partner repository^72^ with accession IDs PXD051311 and PXD051180, respectively. RNA sequencing and ribosome profiling data have been deposited to the GEO database with accession IDs GSE267077 and GSE267078, respectively.

---

**Figure S1.**
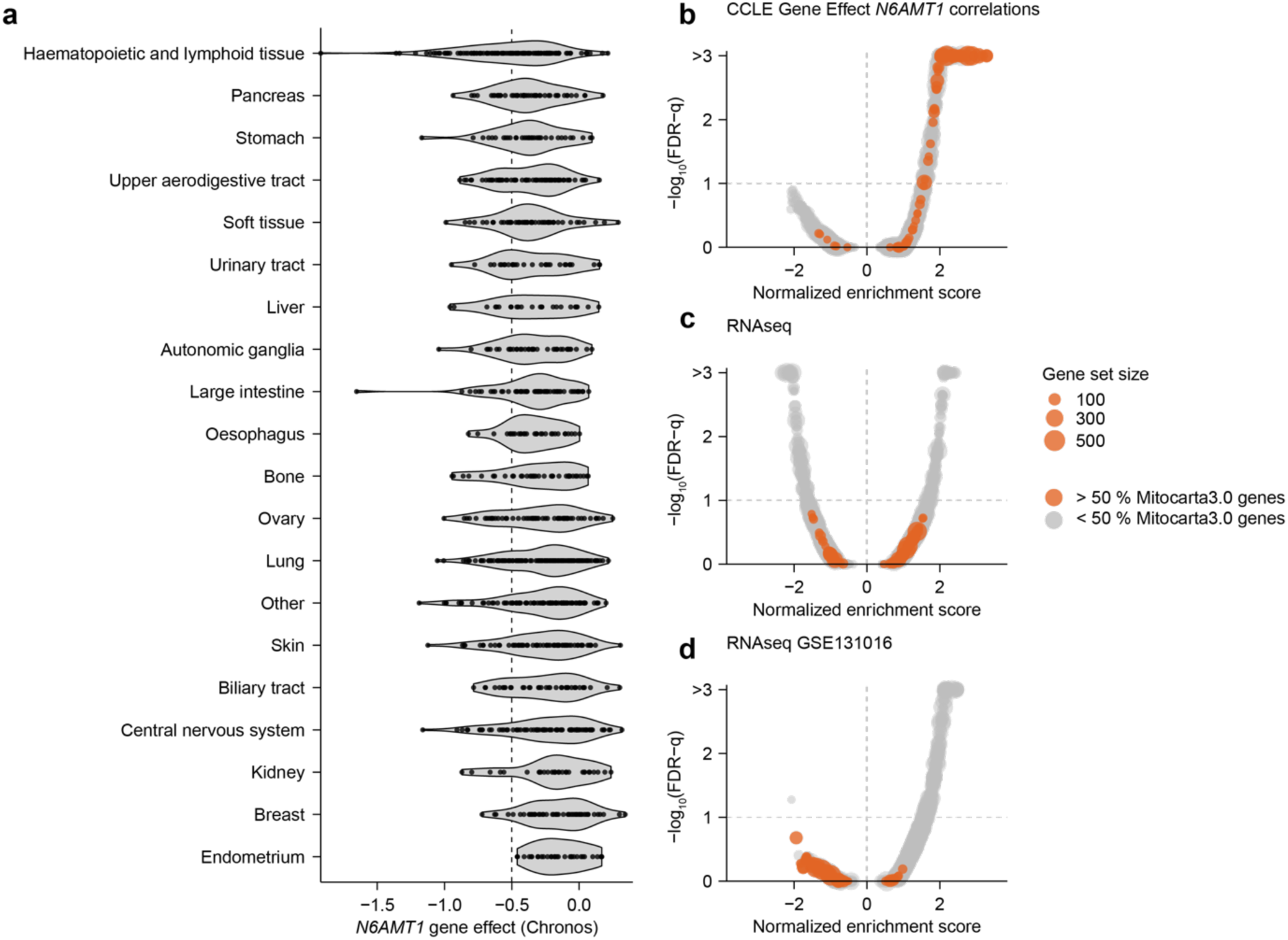
Additional analyses, related to Figure 1. **a.** Violin plots showing distributions of *N6AMT1* CCLE Chronos scores by lineage. Lineages are ordered by increasing mean Chronos score. **b.** Gene set enrichment analyses of correlations shown in Figure 1B. Gene sets with more than 50% MitoCarta3.0 genes are highlighted in orange. **c-d.** Gene set enrichment analyses of RNA-Seq shown in Figure 1F (**c**), and a publicly available RNA-Seq dataset comparing A549 control cell lines with siRNA-mediated *N6AMT1* knockdown (dataset GSE131016 in the Gene Expression Omnibus database) (**d**) showing no change in MitoCarta3.0 transcripts. Gene sets with more than 50% MitoCarta3.0 genes are highlighted in orange.

**Figure S2.**
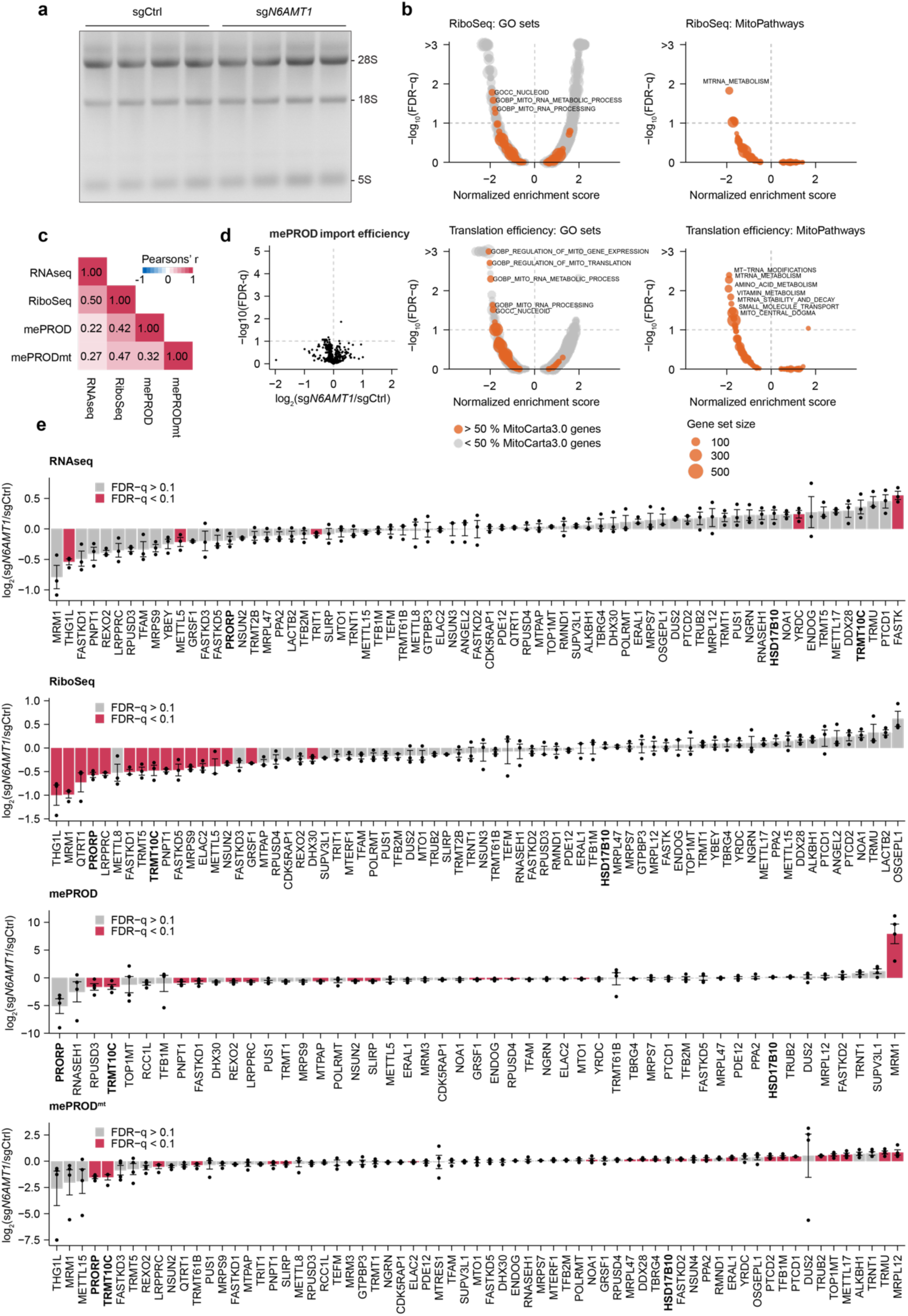
Additional analyses related to Figure 2. **a.** RNA-agarose gel (basis for the quantification in 2A), comparing control to *N6AMT1*-depleted K562 cell lines 10 days post sgRNA transduction (n = 4 independent lentiviral infections per condition). Major ribosomal RNA (rRNA) peaks are indicated. **b.** Gene set enrichment analyses of log_2_(fold-change) values from ribosome profiling analysis (upper panels) and calculated translation efficiencies (ribosome-protected fragments / mRNA abundance; lower panels) based on Gene Ontology (GO) terms (left panels) or MitoPathways gene sets (right panels). Gene sets with > 50% MitoCarta3.0 genes are highlighted in orange. **c.** Pearson’s correlation coefficients between RNA-Seq, Ribo-Seq, mePROD, and mePROD^mt^ datasets. **d.** Mitochondrial import analysis (log_2_-transformed fold-changes in mePRODmt expression / mePROD expression ratios) in *N6AMT1*-depleted K562 as compared to control cells. n = 4 independent lentiviral transduction per condition. **e.** Bar plots showing ranked log_2_(fold-changes) of all genes belonging to the “mt-RNA metabolism” MitoPathways gene set in each of the four indicated datasets. Points represent log_2_(fold-change) values for each independent replicate in the *N6AMT1*-depleted cells as compared to K562 cells. Error bars represent mean +/- standard error of the mean (n = 3-4 independent transduction per condition). Genes highlighted in red are significant at a false-discovery rate (FDR) of 0.1 (Benjamini-Hochberg).

**Figure S3.**
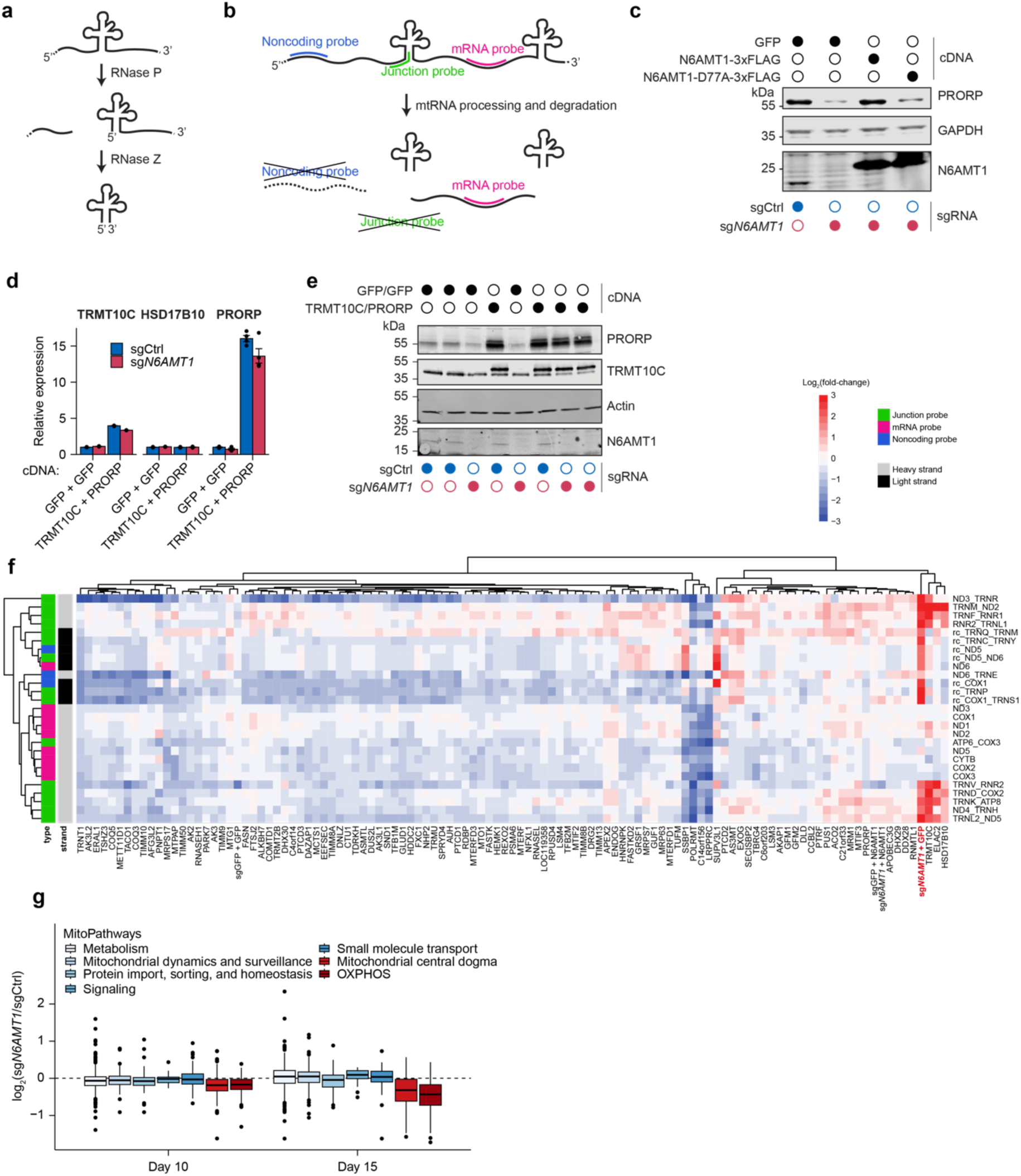
Additional analyses related to Figure 3. **a.** Schematic representation of mitochondrial RNA processing by mitochondrial RNase P and Z. Mitochondrial RNase P is composed of TRMT10C (MRPP1), HSD17B10 (MRPP2), PRORP (MRPP3). **b.** Illustration of MitoString to monitor mt-RNA processing defects. Three classes of probes targeting tRNA-mRNA junctions, noncoding regions, or regions within mRNAs are employed. Upon processing and degradation, junction-targeted probes or probes targeting non-coding regions are unable to bind their corresponding mt-RNA, thus showing reduced signal relative to mRNA-targeted probes. **c.** Protein immunoblot showing mitochondrial RNase P subunit PRORP protein levels in control (sgCtrl) and *N6AMT1*-depleted (sg*N6AMT1*) K562 cells with introduction of cDNA encoding either GFP, wild type N6AMT1 or catalytically inactive N6AMT1-D77A. **d.** qPCR analysis of RNase P-encoding transcripts in control or *N6AMT1*-depleted K562 cells, with introduction of either 2 different *GFP* cDNAs or *TRMT10C* + *PRORP* cDNA (n = 3 independent lentiviral infections). **e.** Immunoblot showing mitochondrial RNase P subunits in control or *N6AMT1*-depleted K562 with the introduction of either 2 different *GFP* cDNAs or a combination of *TRMT10C* and *PRORP* cDNA. **f.** Unsupervised clustering of MitoString data presented in Figure 3A with that of Wolf and Mootha (2014), illustrating the clustering of *N6AMT1*-depletion (sg*N6AMT1* + GFP) with depletion of mitochondrial RNase P (TRMT10C, HSD17B10) and RNase Z (ELAC2) proteins. **g.** Proteomics protein abundance of mitochondrial genes annotated as each of the six top-level pathway annotations in the MitoPathways database.

